# DANGO: Predicting higher-order genetic interactions

**DOI:** 10.1101/2020.11.26.400739

**Authors:** Ruochi Zhang, Mihir Bafna, Jianzhu Ma, Jian Ma

**Affiliations:** Ray and Stephanie Lane Computational Biology Department, School of Computer Science, Carnegie Mellon University, Pittsburgh, PA 15213, USA; MIT Computer Science & Artificial Intelligence Laboratory, Cambridge, MA 02139, USA; Department of Electronic Engineering, Tsinghua University, Beijing, China; Institute for AI Industry Research, Tsinghua University, Beijing, China

## Abstract

Higher-order genetic interactions, which have profound implications for understanding the molecular mechanisms of phenotypic variation, remain poorly characterized. Most studies to date have focused on pairwise interactions, as designing high-throughput experimental screenings for the vast combinatorial search space of higher-order molecular interactions is dauntingly challenging. Here, we develop Dango, a computational method based on a self-attention hypergraph neural network, designed to effectively predict higher-order genetic interaction among groups of genes. As a proof-of-concept, we provide comprehensive predictions for over 400 million trigenic interactions in the yeast *S. cerevisiae*, significantly expanding the quantitative characterization of such interactions. Our results demonstrate that Dango accurately predicts trigenic interactions, uncovering both known and novel biological functions related to cell growth. We further incorporate protein embeddings and model uncertainty scoring to enhance the biological relevance and interpretability of the predicted interactions. Moreover, the predicted interactions can serve as powerful genetic markers for growth response under diverse conditions. Together, Dango enables a more complete map of complex genetic interactions that impinge upon phenotypic diversity.

## Introduction

Genetic interactions describe phenomena where different genes (or more broadly, genetic variants) work synergistically to impact phenotypes [1, 2]. Characterizing the principles and mechanisms of genetic interactions, which have profound implications in understanding human health and disease, has been a central question in systems biology [3–7]. Leveraging the model organism *S. cerevisiae*, both high-throughput experimental approaches and computational methods have been developed to map the landscape of genetic interactions [8–10]. Genetic interactions are often manifested when mutations in the interacting genes lead to unexpected growth phenotypes that cannot be explained by the additive effects of individual mutations [8]. For instance, positive genetic interactions occur when mutations in multiple genes lead to greater growth fitness than expected from individual mutations, while negative genetic interactions, including the extreme form of “synthetic lethality” [11], result in much worse growth fitness than expected. Together, measuring genetic interactions has shed new light on functional dependencies between genes, potential redundancies in the genome, and, crucially, the complex genotype-to-phenotype relationships.

However, most studies on genetic interactions have focused on pairwise, digenic interactions [3, 8]. Growing evidence suggests that higher-order genetic interactions involving three or more genes also play critical roles in controlling phenotypes. Several studies have revealed higher-order genetic interactions across diverse biological contexts [12–18]. For example, by mutating tens of thousands of loci in the *E. coli* genome, a higher-order interaction was identified [15] among Fis, Fnr, and FadR related to cell viability under high acetate concentrations. The “green monster” technology [13] was developed to precisely delete 16 ATP-binding cassette transporters in a drug-sensitive *S. cerevisiae* strain, enabling the delineation of higher-order interactions among these transporters in 5,353 engineered strains with >85,000 genotype-to-resistance drug-response observations [19]. In a landmark study, trigenic interactions in *S. cerevisiae* were systematically profiled [9], determining how different combinations of three-gene deletions impacted colony growth. This study greatly expanded the scope of genetic interaction analysis. However, even this most systematic study of trigenic interactions to date [9] only tested tested ∼100,000 of the possible 36 billion trigenic interactions, leaving the vast majority unexplored.

Here, we develop a new machine learning model, Dango, based on a self-attention hypergraph neural network, to predict higher-order genetic interactions by capturing the complex relationships between higher-order interactions and pairwise interactions extracted from a collection of heterogeneous molecular networks. To evaluate the method, we utilize the trigenic interaction data from [9]. We apply Dango to multiple pairwise interaction networks to predict trigenic interactions. Our results demonstrate that Dango can accurately predict trigenic interactions, outperforming state-of-the-art algorithms generalized from methods designed for pairwise interaction predictions. Notably, Dango not only refines the measured trigenic interactions [9] but also further predicts more than 400 million interaction scores for previously unmeasured trigenic interactions. This significantly expands the repertoire of quantitative trigenic interactions. Furthermore, we show that the trigenic interactions predicted by Dango capture both known and novel functions related to cell growth. When applied to an independent yeast dataset, Dango further demonstrates that predicted trigenic interactions can serve as valuable features to improve mapping from genotype to phenotype.

## Results

### Overall design of Dango

**Fig**. 1a illustrates the overall architecture of Dango for predicting trigenic interactions by modeling genes and their trigenic interactions as a hypergraph. Dango captures higher-order interaction patterns through four main components:

**Figure 1:**
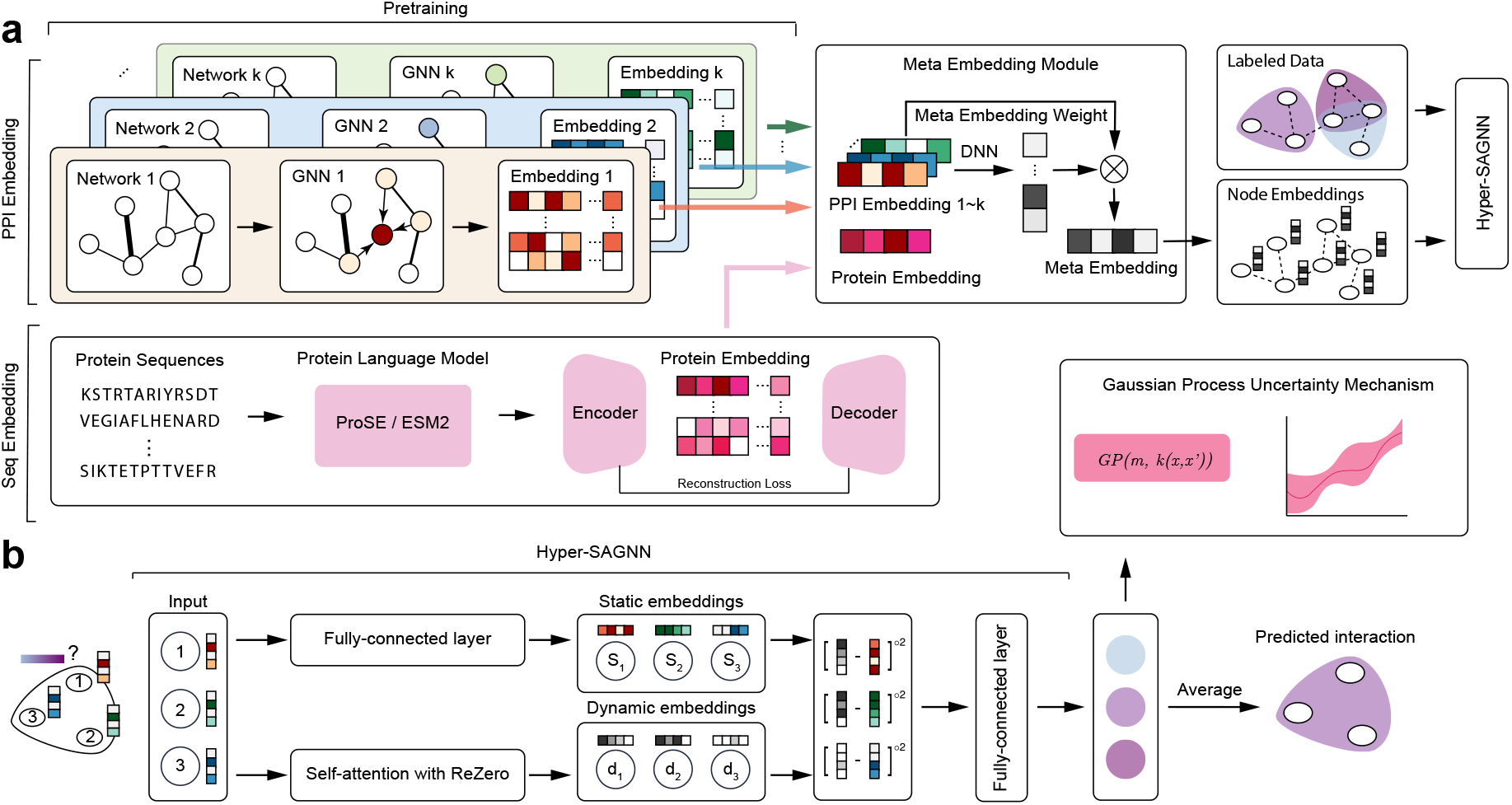
Overview of the Dango algorithm. **a**. Workflow of Dango . Dango takes multiple pairwise molecular interaction networks as input and pre-trains separate graph neural networks (GNNs) to generate node embeddings. An optional protein sequence embedding (ProSE/ESM2) can be included per gene. Dimensionality of the protein embedding is reduced via an autoenoder during pretraining. These embeddings for the same node across different networks are integrated through a meta embedding learning scheme. The integrated node embeddings, along with labeled data trigenic interaction datasets, are used as input to train a hypergraph representation learning framework. **b**. Modified Hyper-SAGNN architecture for hyperedge regression. The model input consists of gene triplets with corresponding node embedding features. These triplets pass through fully-connected layers and modified self-attention layers to generate static and dynamic node embeddings, respectively. A pseudo-Euclidean distance is calculated for each pair of static and dynamic embeddings, and the results are averaged to produce the final predicted trigenic interaction score. An optional Gaussian Process Regressor can be fit to the residuals, bounding each prediction with an uncertainty value (variance).

- Hypergraph construction. Genes are represented as nodes, and trigenic interactions are represented as 3-way hyperedges (marked as “Labeled Data” in **Fig**. 1a). The trigenic interaction scores are modeled as attributes of the hyperedges.
- Pre-training graph neural networks (GNNs). Node embeddings are generated based on six pairwise interaction networks from the STRING database [20].
- Meta embedding learning. Embeddings from different networks are integrated using a meta embedding learning module.
- Hypergraph representation learning. The Hyper-SAGNN [21] model with both labeled data and node features as inputs (**Fig**. 1b). A distinct loss function in Dango is designed to predict hyperedge attributes in a regression manner, different from other applications of Hyper-SAGNN.

Although these components are described separately, the GNNs (after pre-training), the meta embedding learning module, and the Hyper-SAGNN are optimized jointly in an end-to-end manner. Importantly, Dango is generalizable to higher-order interactions beyond trigenic ones. Detailed structures of these components are described in **Methods**. Additional details on model choice, training procedure, computational complexity, and the baseline model designs are provided in the **Supplementary Methods**.

### Datasets used in Dango

The six PPI networks used as input to the GNNs are derived from STRING database v9.1 [20]. Although the latest STRING database v11.5 contains more PPIs, we used v9.1 to ensure fair comparisons with baseline models. These networks are categorized as: “Experimental” (experimentally verified PPIs through literature curation), “Database” (PPIs imported from other database), “Neighborhood” (proteins with similar genomic context in different species), “Fusion” (proteins fused within a given genome), “Co-occurrence” (proteins with similar phylogenetic profile), and “Co-expression” (predicted association based on co-expression patterns).

The trigenic interaction dataset analyzed in this work comes from [9], comprising ∼91,000 measured trigenic interaction scores (τ) across 1,400 genes. Of these, 1,395 genes also appear in the pairwise PPI networks used as input, and triplets involving the five excluded genes were removed from further analysis. Notably, all measured trigenic interaction scores in this dataset are negative, with no observed positive trigenic interactions. The Pearson correlation between the trigenic interaction scores from two individual replicates is approximately 0.59, which is much lower than the correlation observed for digenic interaction scores (0.88) from the same data source. These values represent an approximate upper bound for achievable computational performance, highlighting the challenges in modeling higher-order interactions from the dataset.

### Dango accurately predicts trigenic interactions

We systematically evaluated Dango ’s performance in predicting trigenic interaction scores and compared it against three baseline models: Mashup-RF, Mashup-GTB (based on the state-of-the-art biological network integration method Mashup [22]) and GCN-Avg-DNN (a graph neural network model trained in an end-to-end manner). Details on training procedure, computational complexity, and baseline model designs are provided in **Supplementary Methods**. Prediction performance was assessed using the following metrics: (1) Pearson and Spearman correlations between predicted and measured trigenic interactions; (2) AUROC and AUPR for classifying triplets into “strong” and “weak” interactions with a cutoff 0.05 on the absolute measured interaction score; (3) Pearson and Spearman correlations within the “strong” interaction subset.

Using 5-fold cross-validation on the trigenic interaction dataset [9], we collected predictions for all test sets across folds. Dango demonstrated accurate predictions for trigenic interactions (**Fig**. 2a) and significantly outperformed all baseline models across all evaluation metrics (**Fig**. 2b). Among the baselines, GCN-Avg-DNN showed better performance than Mashup-RF and Mashup-GTB, suggesting that end-to-end model training allows node embeddings to more effectively capture task-specific information.

**Figure 2:**
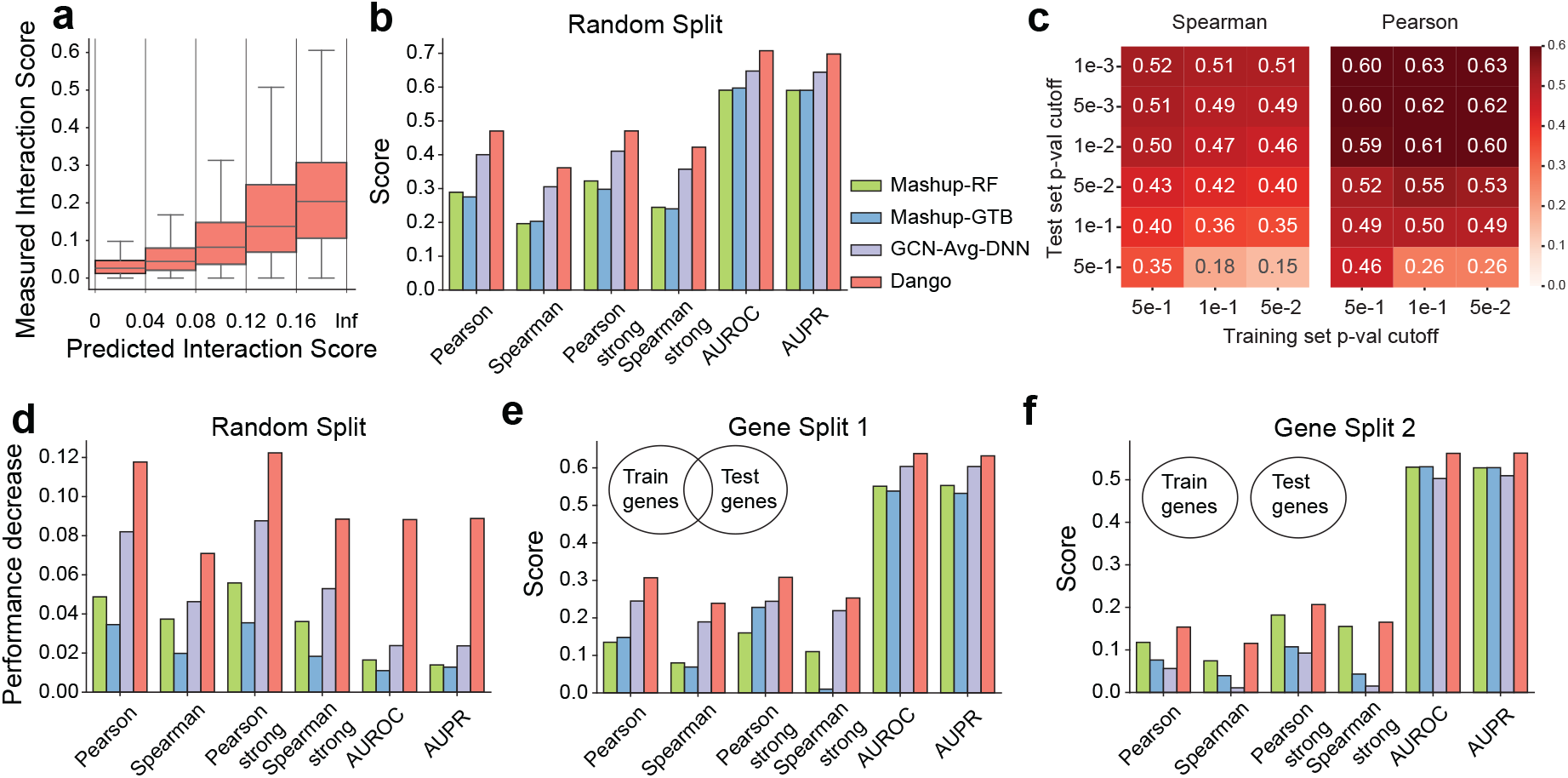
Evaluation of performance on trigenic interaction score predictions. **a**. Comparison of measured versus predicted trigenic interaction scores for each triplet gene disruption genotype. **b**. Evaluation of the predicted trigenic interaction scores with random partition cross-validation. Metrics include Pearson and Spearman correlation scores, Pearson and Spearman correlation scores within strong trigenic interactions ((|τ| > 0.05)), and AUROC/AUPR scores for predicting whether a measured interaction score is strong or not. **c**. Analysis of performance changes when using different *P*-value cutoffs on the training and test datasets. *P*-values reflect the consistency of trigenic interaction scores calculated on two replicates. **d**. Decrease in performance when the learned gene embeddings are randomly shuffled. For GNN-based methods, the adjacency matrix of the input network is shuffled. Evaluations are conducted under random cross-validation. **e, f**. Evaluation of performance under two gene-based training and testing set split. In Gene Split 1, around 60 genes in the test set are unobserved in the training data. In Gene Split 2, all 400 genes in the test set are unobserved in the training data.

The trigenic interaction dataset includes replicate measurements, enabling calculation of a *P*-value for consistency between replicates. We tested whether Dango would achieve higher correlation scores on a smaller, more reliable subsets of data filtered by *P*-value thresholds. As shown in **Fig**. 2c, Pearson and Spearman correlations improve as the *P*-value cut-off becomes stricter, particularly between 0.5 and 0.01. Crucially, Dango maintains strong performance on these reliable test sets even with less reliable training data. Of the measured ∼100,000 trigenic interaction scores [9], only 4,042 pass the stringent *P*-value cut-off 0.01 (see **Fig**. S3), underscoring the difficulty in obtaining reliable scores for all gene combinations. Dango ’s stable performance highlights its ability to uncover meaningful interactions from noisy data and refine the trigenic interaction scores originally measured by experimental approaches.

### Assessing bias in trigenic interaction predictions

We sought to assess whether the performance improvements of Dango were due to its enhanced ability to capture interaction patterns or simply “memorizing” potential biases (i.e., overfitting to confounding factors present in both training and test sets) [23]. For examples, if a gene frequently appears in the measured trigenic interaction dataset with high interaction scores, the model could achieve good performance simply by assigning high scores to all triplets containing that gene. To evaluate this, we specifically tested for the presence of such bias in Dango and the baseline models using two approaches.

First, we randomly shuffled the node embeddings and retrained the model. In this setup, each gene was assigned the embedding vector of another gene. For Mashup-RF and Mashup-GTB, this was achieved by shuffling Mashup embeddings across genes. For GCN-Avg-DNN and Dango, we shuffled the adjacency matrix of the six PPI networks used in the GCN. If the model’s strong performance were solely due to overfitting to the training set, random shuffling would not lead to a significant performance drop. Conversely, if the model learns meaningful interaction patterns, shuffling would cause a noticeable decrease in performance. Among all methods, Dango exhibited the most significant performance reduction when features were shuffled across genes (**Fig**. 2d), while the baseline models showed much smaller decrease, indicating a greater susceptibility to potential bias from the models.

Second, we used alternative training/test split schemes to evaluate whether the models could generalize to unseen genes. In the first scheme, we excluded around 60 genes from the training set, ensuring that all triplets containing at least one of these 60 genes were included included in the test set (**Fig**. 2e). In the second scheme, we selected 400 genes and included only triplets involving three genes from this set in the test set, while excluding triplets with one or two of these genes from both training and test sets (**Fig**. 2f). In other words, we focus on evaluating triplets with three unobserved genes. Under both settings, Dango consistently achieved the best performance among all methods (**Fig**. 2e-f). Additionally, we evaluated performance decrease after shuffling features under these two training/test split schemes. Consistent with the observations from the random split scheme (**Fig**. S2), Dango again exhibited the largest performance reduction for almost all metrics.

These evaluations collectively suggest that Dango can accurately predict trigenic interaction scores and its strong performance on the test set is not due to overfitting to confounding factors in the data set. One key reason for Dango ’s ability to overcome such bias is the unique self-attention mechanism in Hyper-SAGNN, which enables meaningful interactions between node embeddings – a feature not guaranteed in the baseline methods.

### Dango discovers novel trigenic interactions with important biological functions

We next investigated whether Dango could be used to make *de novo* predictions of trigenic interactions. We applied the trained Dango model to predict interactions in two sets of gene triplets. The first set consisted of the same triplets measured in [9]. Predictions for this set were obtained through a 5-fold cross-validation, where the test-set predictions from each fold were aggregated. The second set included all possible combinations of the 1,395 genes in the dataset (∼451 million triplets), excluding those already measured in [9]. Predictions for this set were made using a Dango model trained on the entire set of measured trigenic interactions.

For both predicted and measured interactions, we selected the top 20% based on interaction scores for further analysis (with a cutoff of |τ| ∼0.05 for measured trigenic interactions). We characterized the functional properties of these selected interactions through Gene Ontology (GO) enrichment analysis for biological process and cellular component annotations. Genes were assigned to the “leaf node” of the GO hierarchy if they had multiple annotations. Each trigenic interaction was classified as “across terms” or “within terms” based on whether all three genes belong to the same GO term. We calculated fold-changes in the frequency of GO terms in the selected trigenic interactions compared to the background (all triplets in the two sets) and assessed enrichment significance using a hypergeometric test. As shown in **Fig**. 3a-b, when evaluating on set #1, the enriched GO terms largely overlapped with those from the original measured interactions, and their fold-change scores correlated well (Pearson correlation 0.56 for “within terms” and 0.94 for “across terms”). This indicates that Dango effectively captures higher-order genetic interactions from the original datasets. Similarly, in set #2, fold-change scores for GO term patterns present in the original dataset showed strong correlations (Pearson correlation 0.53 for “within terms” and 0.72 for “across terms”). However, the predicted interactions in set #2 also revealed enrichment for additional GO terms, many of which were associated with cell growth. Notably, cellular components such as ‘cytoplasmic vesicle’, ‘cell cortex’, and ‘cytoskeleton’ – critical for cell survival – were enriched. Additional processes, including ‘RNA splicing’, ‘Golgi vesicle transport’, ‘Golgi apparatus’, and ‘microtubule organizing center’, are also known to play vital roles in regulating growth and development [24–26].

**Figure 3:**
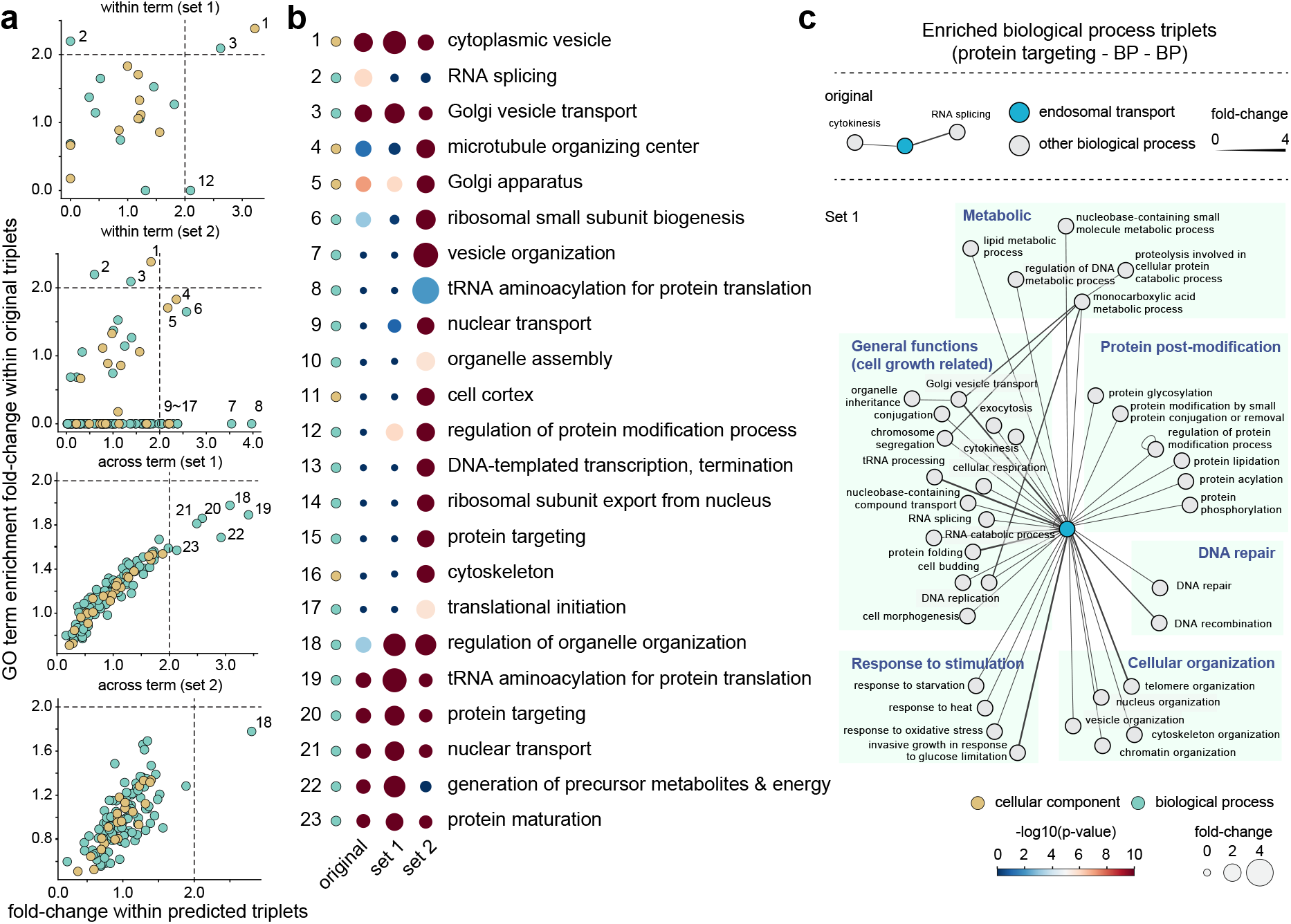
Functional characterization of the predicted trigenic interactions. **a**. Correlation of frequency fold-change of trigenic interactions within or across Gene Ontology (GO) terms between predicted and measured trigenic interactions. Two sets of predicted trigenic interactions are analyzed: set #1 corresponds to the same triplets measured in [9], and set #2 corresponds to all possible combinations of three genes that appear in [9]. **b**. Enriched GO terms across different set of trigenic interactions. GO terms with a frequency fold-change greater than 2.0 in any set are visualized, showing the frequency fold-change and associated *P*-value (hypergeometric test). **c**. Discovery of a functionally important trigenic interaction hub by Dango . Networks visualize enriched combinations of biological process terms, including a novel trigenic interactions hub identified in set #1 that was not observed in the original dataset. This hub comprises genes related to protein targeting, endosomal transport, and other biological process (e.g., protein post-modification, DNA repair).

We further analyzed combinations of different GO terms by calculating fold-changes for their co-occurrence in selected trigenic interactions compared to the background. Dango uncovered significant combinations of biological processes that were not enriched in the original dataset. For instance, **Fig**. 3c highlights a novel trigenic interaction hub involving genes related to protein targeting and endosomal transport (*P*-value ≤ 10^−3^). Within the network for set #1, the node corresponding to the biological process ‘endosomal transport’ had a high degree, reflecting a hub of trigenic interactions linking genes related to protein targeting, endosomal transport, and various other biological processes.

We also examined whether the identified interactions co-occurred frequently in higher-order interactions captured by orthogonal data, such as KEGG pathways [27] We calculated fold-changes for the co-occurrence of the three genes in the trigenic interaction within specific KEGG pathways compared to the background (**Fig**. 4a). In the original dataset, only two KEGG pathways (MAPK signaling pathway and cell cycle) significantly overlapped with the predicted trigenic interactions (*P*-value ≤ 10^−3^, hyper-geometric test). Both pathways had lower *P*-values in set #1, which also revealed four additional trigenic interactions involving these pathways. Notably, two of these interactions (BEM3, RGA1) had high predicted interaction scores and recorded physical interactions with CLN1/2 based on BioGRID [28]. Set #1 also uncovered an additional significant KEGG pathway, endocytosis, containing three triplets with all genes from this pathway and highly ranked predicted scores. For set #2, Dango has identified all these KEGG pathways with much higher fold-change value and lower *P*-values, but it further revealed 14 additional KEGG pathways with significant enrichment.

**Figure 4:**
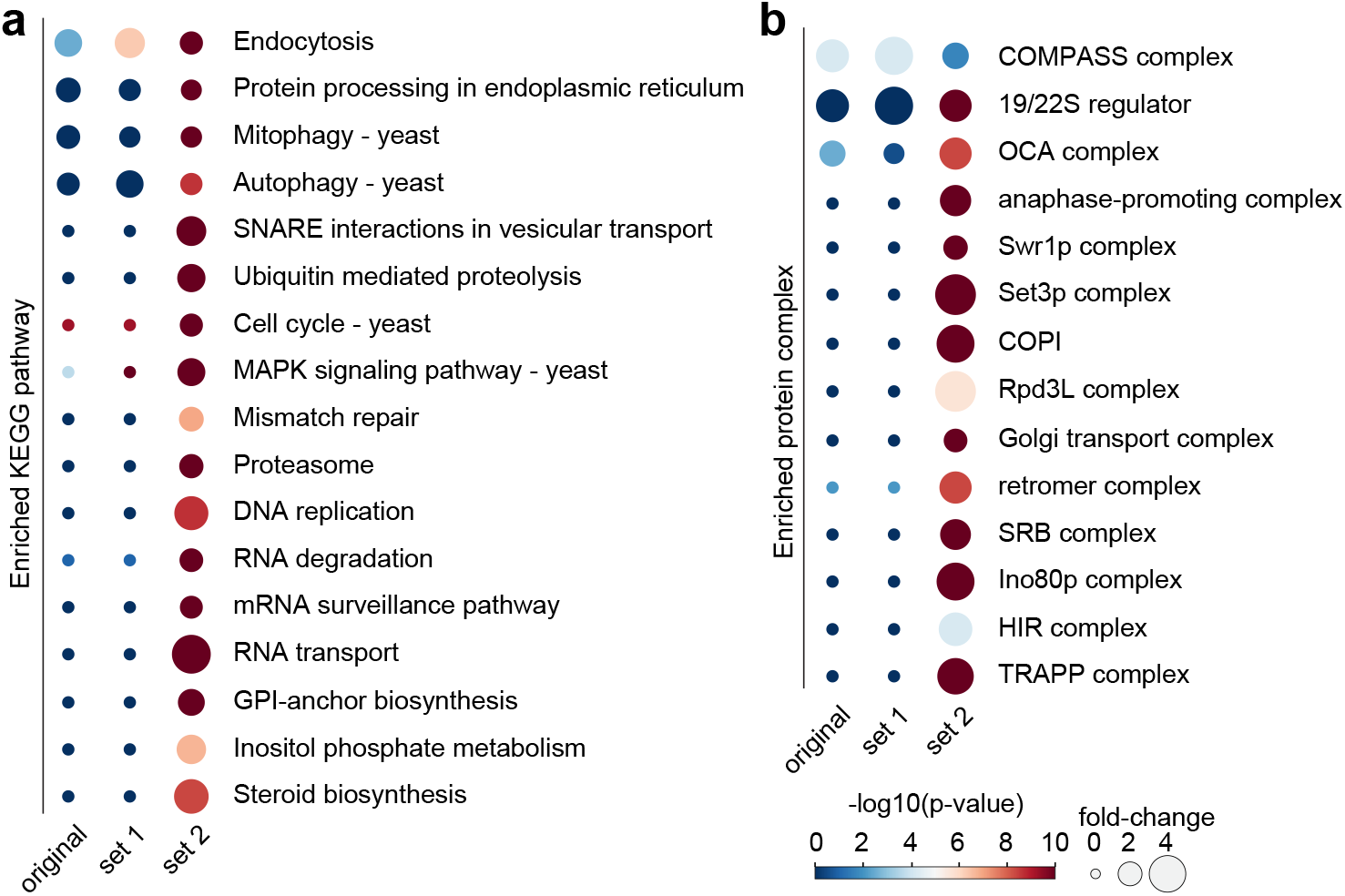
Enrichment analysis of KEGG pathways and yeast complexes in predicted trigenic interactions. **a**. Enriched KEGG pathways across different sets of trigenic interactions. Pathways are visualized with their frequency fold-change and *P*-values calculated using the hypergeometric test. **b**. Enriched yeast protein complexes across different sets of trigenic interactions. Complexes are visualized similarly, with their frequency fold-change and hypergeometric test *P*-values.

We further tested whether the predicted interactions were enriched for known protein complexes by comparing them to a curated protein complex database [29] (**Fig**. 4b). In both the original trigenic interaction dataset and set #1, only one complex (COMPASS complex) showed significant overlap with the trigenic interactions. In contrast, set #2 revealed 14 additional protein complexes with significant enrichment.

These findings demonstrate that the trigenic interactions predicted by Dango can uncover both known and novel biological functions related to cell growth, highlighting its potential to facilitate the discovery of new biological pathways and protein complexes.

### Trigenic interactions predicted by Dango enhance the prediction of yeast growth

To further demonstrate that the trigenic interactions predicted by Dango provide critical insights into the wiring mechanism of gene functions and pathways, we evaluated its utility in predicting yeast growth under various conditions that affect physiological and cellular responses of distinct yeast strains. Growth data was obtained from [30], which characterized the single nucleotide polymorphisms (SNPs) of 971 yeast strains and quantified their growth fitness phenotype under 35 conditions (e.g., caffeine addition or high-temperature culturing). Our goal was to use the SNPs of each strain to predict its 35 growth fitness scores. This prediction task is highly challenging due to the high dimensionality of the data: the dataset contains over 18,000 SNPs, exceeding the number of yeast strains available for training. Even when SNPs are grouped by the genes they reside in, the feature dimension for each strain exceeds 6,000. The high dimensionality, combined with insufficient data points, make purely data-driven approaches, such as deep neural networks (DNNs), prone to overfitting or converging to less preferred local optima.

To address this, we incorporated the identified trigenic interactions as priors into the neural network structure, enforcing them to improve the prediction of growth fitness scores (see **Fig**. 5a for model structure; **Supplementary Methods** for detailed descriptions). Depending on whether SNPs were grouped by genes as input to the model, we referred to these models as Sparse (trigenic)-gene and Sparse (trigenic)-SNP, respectively. For fair comparisons, we limited SNPs to the 1,395 genes included in [9]. As baseline methods, we implemented a two-layer neural network with fully connected layers, referred to as Dense-gene and Dense-SNP, respectively. The hidden layer dimension was tuned between 8 to 1,024 and optimized at 256.

**Figure 5:**
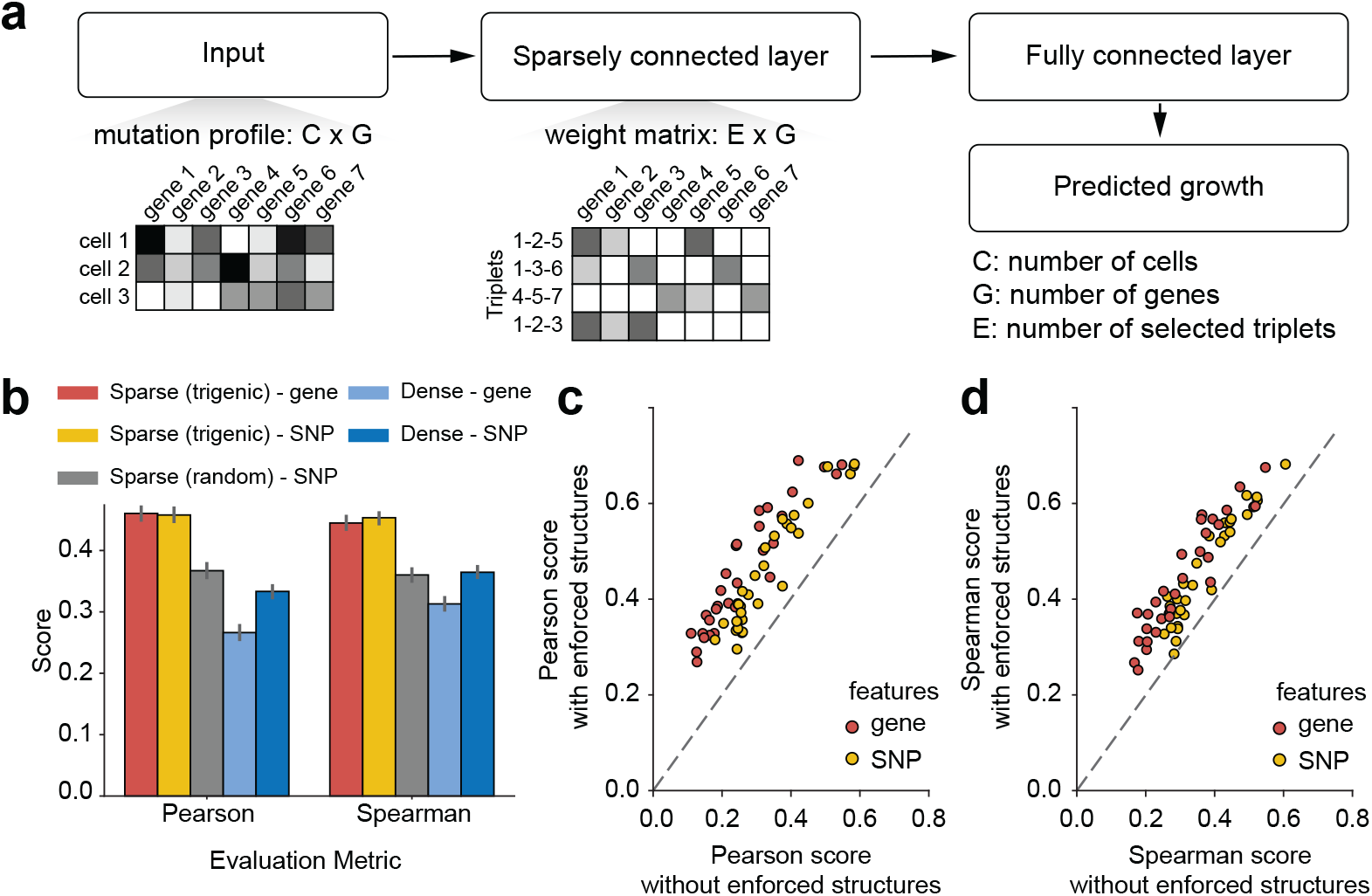
Trigenic interactions identified by Dango predict yeast growth under unseen culture conditions. **a**. Illustration of the integration of selected trigenic interactions into the structure of a deep neural network (DNN). The selected interactions guide the network architecture to improve prediction accuracy. **b**. Performance evaluation of trigenic interaction-guided DNNs compared to baseline models for predicting yeast growth. The performances are aggregated across all 35 conditions. **c, d**. Performance evaluation of trigenic interaction-guided DNNs versus baseline models under each individual condition.

As shown in **Fig**. 5b, each model was trained and evaluated 10 times. Across all conditions, models with enforced structures based on trigenic interactions (Sparse (trigenic)-SNP and Sparse (trigenic)-gene) consistently outperformed DNNs without sparsity constraints (Dense-SNP and Dense-gene), regardless of whether raw SNP frequencies or gene-aggregated SNPs were used as input. Comparing Pearson and Spearman correlations under each condition, the performance improvement of models guided by trigenic interactions was robust across all conditions (**Fig**. 5c-d). To determine whether the improvement resulted from prior knowledge of trigenic interactions or merely from sparsity constraints reducing overfitting, we constructed a baseline model with the same sparse weight matrix structure as Sparse (trigenic)-SNP but with a random structure. This model, Sparse (random)-SNP, retained the same sparsity ratio as the trigenic interaction-guided model. Importantly, the Sparse (trigenic)-SNP model outperformed the Sparse (random)-SNP model (**Fig**. 5b). Additionally, the Sparse (random)-SNP model performed similarly to the Dense-SNP model, suggesting that the improvement observed in Sparse (trigenic) models was indeed due to the incorporation of trigenic interaction knowledge rather than the sparsity constraint alone.

These results demonstrate that trigenic interactions identified by Dango can serve as effective prior for predicting growth fitness under various conditions. They also provide strong evidence that Dango can transfer the model trained under one condition to other settings, offering potential for advancing our understanding of complex genotype-to-phenotype relationships across diverse biological contexts.

### Protein embeddings and model uncertainty scoring in Dango reveals novel higher-order interactions

We sought to further enhance Dango by incorporating protein embeddings and introducing model uncertainty scoring to refine trigenic interaction predictions. Language models [31, 32] have demonstrated the ability to embed biological sequences into representations that preserve properties of sequence, structure, and function. To improve Dango ’s trigenic interaction predictions, we incorporated protein embeddings for yeast genes to provide additional insights into their structure and function. These embeddings were incorporated during the Dango pretraining step, where an autoencoder was used to reduce their dimensionality to match the pretrained embeddings from other sources (combined via the MetaEmbedding module). Details of the protein autoencoder are described in **Methods**.

To provide explainability for Dango ’s predictions, we adopted a Gaussian process (GP) regression framework for uncertainty scoring [33, 34]. The GP model was trained on residuals between predicted and true trigenic interaction scores, outputting both a mean prediction (which improves accuracy when combined with the original Dango score) and a variance (representing the model’s uncertainty in each prediction). Further details on the GP formulation are provided in **Methods.** **Fig**. 6a highlights a general trend where Dango exhibits higher uncertainty for interactions with higher predicted scores, likely because such high-scoring interactions are rarer. This observation underscores the significance of identifying interactions that are both high-scoring and low in uncertainty.

We refined trigenic interaction scores using the GP uncertainty mechanism by ranking interactions based on high Dango scores and low uncertainty, then combining these ranks via the harmonic mean to prioritize high-confidence, high-scoring interactions. This approach yielded an even stronger association between predicted trigenic interactions and protein complexes (**Fig**. 6b), validating both the inclusion of protein embeddings and the use of uncertainty scoring to enhance prediction and explainability.

**Figure 6:**
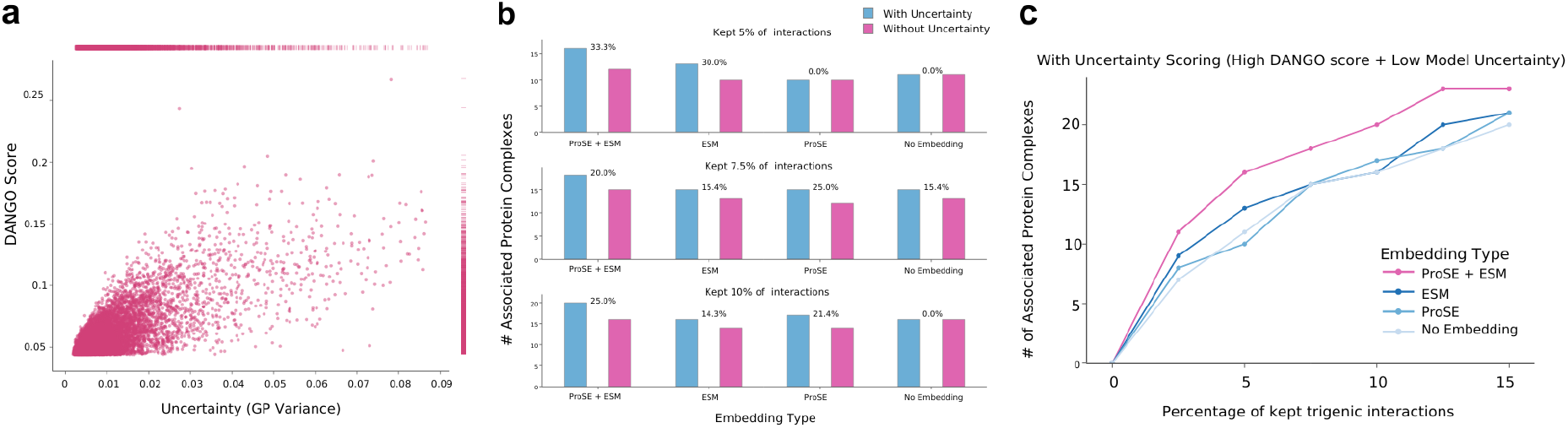
Relationship between Dango scores, model uncertainty, and protein complex associations **a**. Scatter plot of 10,000 random predicted trigenic interactions highlighting the Dango score vs. the model uncertainty. As Dango score increases, model uncertainty tends to increase as well. Panel **b**-**c** show the effect of protein embeddings (ProSE, ESM2) on the number of trigenic interactions associated with protein complex formation across various score thresholds. **b**. Number of associated protein complexes when using and without using the uncertainty score combined with the original high Dango scoring trigenic interactions. The change in number of associated protein complexes after using uncertainty scores is expressed as percentage. **c**. Number of associated protein complexes using a combination of high Dango scores and low model uncertainty, calculated as the harmonic mean of the two metrics. Including model uncertainty results in increased protein complex associations.

We hypothesized that incorporating protein embeddings would improve Dango ’s ability to identify trigenic interactions associated with physically interacting protein complexes. To test this, we used protein embeddings generated from ProSE [31] and ESM2 [32]. For each gene, we generated trigenic interaction scores using Dango where the pretraining included the protein embeddings. Across thresholds of top-scoring interactions (ranging from 0 to 20%, increasing in increments of 2.5%), the proportion of trigenic interactions associated with known protein complexes consistently increased when protein embeddings were included (**Fig**. 6c).

To properly assess the effectiveness of including these protein embeddings, we compared three models: one pretrained with ProSE embeddings, another with ESM2 embeddings, and a control model without embeddings. High-scoring, low-uncertainty interactions unique to the embedding-guided models (for both ProSE and ESM2) revealed significant association with the yeast SAGA complex, a key regulator for transcription through its histone acetylation function [35]. The complex, resolved via Cryo-EM into 19 subunits organized into four submodules (HAT, DUB, TAF, SPT) [36], showed strong trigenic interactions within submodules (e.g., Spt3, Spt7, Spt8 in SPT; Taf12 in TAF) and across submodules (visualized in **Fig**. S5a). Additionally, interactions were identified within the SLIK complex, a SAGA-like complex with the unique component RTG2 [37], which was also predicted by Dango (**Fig**. S4b).

These findings suggest that protein embeddings enable Dango to identify trigenic interactions linked to physical complex formation, distinguishing them from interactions related to transient signaling pathways. The structural and sequence information inherent in these embeddings provides valuable context for trigenic interaction predictions.

Together, these evaluations highlight the utility of additional embeddings, such as protein embeddings, in guiding Dango without sacrificing prediction accuracy. Furthermore, the GP uncertainty mechanism provides a systematic approach to assess and explain model predictions, offering a confidence score for each predicted interaction.

## Discussion

In this work, we developed Dango, the first self-attention hypergraph neural network designed to model and predict higher-order genetic interactions. Dango outperforms existing methods and effectively identifies biologically meaningful trigenic interactions, with enhanced interpretability through protein embeddings and uncertainty modeling. Beyond primary predictions, Dango also generated a large set of novel trigenic interactions, greatly expanding candidates for experimental validation. While this study focused on yeast, Dango is broadly applicable to other species and biological contexts, offering a powerful tool for studying complex genetic interactions [38, 39].

A potential extension is to expand the scope of phenotypic traits modeled by Dango . While this work focused on yeast growth fitness measured by colony size, our results on predicting phenotypes under unseen conditions suggest the potential to apply Dango to other phenotypes, including human disease traits. Although current datasets on perturbations in human cells exist [40], their scale remains limited. Leveraging extensive higher-order interaction data from yeast, combined with human protein homology [41, 42], could enable the prediction of higher-order genetic and perturbation interactions in humans. This approach has the potential to guide experimental designs for studying complex perturbations and inform therapeutic strategies targeting gene networks in human diseases.

Additionally, a method like Dango supports “AI-in-the-loop” and “lab-in-the-loop” paradigms [38], where iterative cycles of computational predictions and targeted experimental validations create a powerful feedback mechanism. For instance, systematically prioritizing candidate interactions based on high predicted scores and low model uncertainty allows laboratories to allocate resources more effectively and uncover novel biological insights without exhaustive experimentation.

## Method

### Definitions

#### Definition 1. (Digenic/Trigenic interactions)

Higher-order genetic interaction are quantified by the differences between observed phenotypes from the joint deletion of multiple genes and the expected phenotypes from single-gene deletions [9]. For the yeast data studied, the phenotype of a strain (cells of the same genotype) refers to growth fitness, with colony size commonly used as a proxy. The *digenic interaction score* (pairwise interaction) *ϵ*_*ij*_ between mutants *i* and *j* is defined as: *ϵ*_*ij*_ = *f*_*ij*_ −(*f*_*i*_*f*_*j*_), where *f*_*ij*_, *f*_*i*_, *f*_*j*_ represent the measured growth fitness of strains with double or single mutations (*i* and/or *j*). Similarly, the *trigenic interaction* score τ is defined as [9]: τ_*ijk*_ = *f*_*ijk*_ − *f*_*i*_*f*_*j*_*f*_*k*_ − *ϵ*_*jk*_*f*_*i*_ − *ϵ*_*ik*_*f*_*j*_ − *ϵ*_*ij*_*f*_*k*_. The definition accounts for both the expected fitness from single mutations and the contributions of lower-order interactions.

#### Definition 2. (Hypergraph)

A *hypergraph* is formally defined as *G* = (*V, E*), where *V* = *{v*_1_, …, *v*_*n*_*}* represents the set of nodes, and 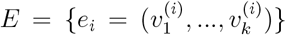 represents the set of hyperedges [43]. A *hyperedge e* connects two or more nodes (|*e*| ≥ 2).

#### Definition 3. (The hyperedge prediction problem)

A key challenges in hypergraph representation learning is the *hyperedge prediction problem* [21, 43–46], which aims to predict the probability of a group of nodes (*v*_1_, *v*_2_, …, *v*_*k*_) forming a hyperedge or to predict the attributes of the hyperedge based on the features of these nodes (*E*_1_, *E*_2_, …, *E*_*k*_).

### GNN pre-training for generating node embeddings

We use six different types of protein-protein interaction (PPI) networks from the STRING database [20] for *S. cerevisiae*, generated based on high-throughput interaction assays, curated PPI databases, and co-expression data. Each network comprises *G* nodes (genes) with weighted edges representing specific types of interactions between the corresponding gene pairs. The following steps are performed on each network independently but in parallel.

For each of the six networks, a two-layered GNN [47] is pre-trained to reconstruct the graph structures. Specifically, for a node *i*, its initial feature vector 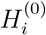 is obtained from a shared embedding layer across all six GNNs. The weight of the embedding layer *H*^(0)^ has size of *G* × *D*, where *G* is the number of genes and *D* is the feature dimension. For each layer in the GNN, the output vector for node *i* aggregates input vectors from its neighbors **𝒩**_*i*_:

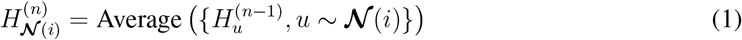

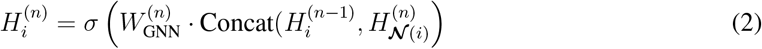

where 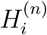 is the output vector of the node *i* at the *n*-th layer of the GNN, 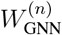 is the weight matrix to be optimized at the *n*-th layer, and *σ* is a non-linear activation function.

The output of each node from the last layer of the GNN (node embeddings) passes through a fully connected layer to reconstruct the corresponding rows in the original adjacency matrix. The loss term *L*_*i*_ for node *i* is:

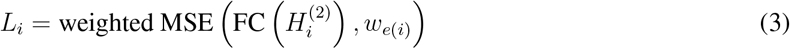

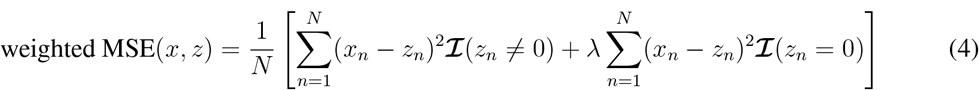

where FC represents the fully connected layer, and *w*_*e*(*i*)_ is the *i*-th row of the network adjacency matrix. *z*_*n*_ is the *n*-th element of the vector *z* of size *N* . The weighted mean-squared error (MSE) function penalizes the zero entries and non-zero entries in the ground truth differently, allowing for nuanced reconstruction of the adjacency matrix. We opted for this weighted MSE over the standard MSE as the reconstruction loss because a zero in the molecular interaction networks does not necessarily imply no interaction between a pair of genes. The hyperparameter *λ* determines the degree to which zeros in the network are considered true non-interactions. To set *λ*, we calculated the percentage of zero entries in the STRING database v9.1 that were confirmed as interactions in the updated v11.0. This percentage, which ranges from 0.02% (co-occurrence) to 2.42% (co-expression), reflects the potential for validation of zero entries in the future. Based on this metric, if the decrease in zeros was greater than 1%, we set *λ* = 0.1; otherwise, we set *λ* = 1.0.

### Optional protein sequence and structure node embedding

Recent advancements in protein language models, such as ProSE [31] and ESM2 [32], have greatly improved performance on sequence and structure-related tasks. To leverage these developments, we incorporated corresponding protein sequence and structure embeddings to potentially enrich the predictions of meaningful trigenic interactions, particularly those involving physical interactions (i.e., protein complex formation). The pretrained ProSE and ESM2 models are used to generate embeddings for each of the *G* nodes corresponding to yeast genes. To ensure compatibility with the GNN pre-trained embeddings, the protein embeddings *X*_*p*_ are reduced in dimensionality using a fully connected autoencoder. Each encoding and decoding layer *i* is denoted as:

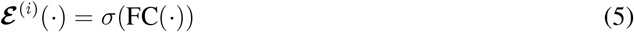

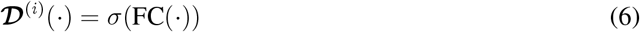

where *σ* represents nonlinear ReLU activation function.

The encoder and decoder each consist of three fully connected layers (FC) with nonlinear ReLU activations. The original protein embeddings *X*_*p*_ ∈ ℝ^*G×O*^ are encoded into lower dimensional representations *Z*_*p*_ ∈ ℝ^*G×D*^ and subsequently reconstructed as 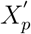 by the decoder. Here, *O* represents the original embedding dimension and *D* represents the reduced dimension, which matches the PPI embeddings. This process is formulated as:

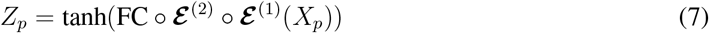

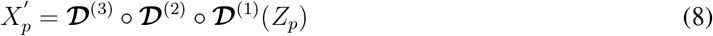

During pre-training, the canonical MSE loss 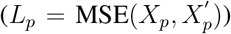 is added to the total loss of the GNN pre-training process described previously.

### Meta embedding learning for the integration of multiple embeddings

The six pre-trained GNNs described in the previous section generate node embeddings that capture the neighborhood topology in each PPI network. Additionally, for each gene, we have optional protein embeddings that capture complex information about the yeast protein’s sequence and structure, which may aid in specific trigenic interaction prediction scenario. In Dango, we aggregate these various embeddings using a meta embedding learning module designed to integrate embeddings of the same node across different networks.

For a node *i*, its node embeddings (six from PPI networks and a seventh optional protein embedding) are denoted as 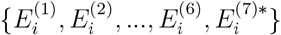. The meta embedding for node *i* is calculated as:

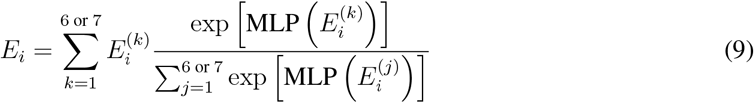

where MLP represents the multi-layer perceptron consisting of two fully-connected layers.

Compared to simpler methods such as averaging or concatenation, meta embedding learning with trainable weights addresses challenges such as unseen entities or inconsistent embedding qualities across different sources [48–50]. For instance, some PPI networks might have no observed interactions for specific genes, but this absence does not necessarily indicate a lack of interaction. Averaging or concatenation would give such embedding equal weight, potentially diluting more informative embeddings. Additionally, the most relevant PPI properties for predicting trigenic interactions may vary across genes. For example, trigenic interactions may arise from functional interchangeability among genes or the formation of protein complexes. Including protein embeddings enables structural information to be incorporated, capturing correlations with protein complex formation. The trainable weights in the meta embedding module allow Dango to adaptively emphasize embeddings based on their relevance.

### Hypergraph representation learning to predict trigenic interactions

Here, we briefly introduce the structure of Hyper-SAGNN (**Fig**. 1b; more details in Ref. [21]), with modifications to improve performance. Each input consists of meta embeddings for three genes, (*E*_1_, *E*_2_, *E*_3_). These embeddings pass through a shared feed-forward neural network, producing static embeddings (*s*_1_, *s*_2_, *s*_3_), where *s*_*i*_ = *σ* (FC (*E*_*i*_)). The static embedding *s*_*i*_ remains constant for a gene *i* across all triplets.

The same input also passes through two multi-head self-attention layers [51], producing dynamic embeddings (*d*_1_, *d*_2_, *d*_3_). Each dynamic embedding *d*_*i*_ depends on all features within the triplet, making it variable across triplets. We denote the weight matrices within each self-attention layer as *W*_*Q*_, *W*_*K*_, *W*_*V*_, which represent the linear transformation of features before applying the scaled dot-product attention [51].

The dynamic embeddings *d*_*i*_ are computed as:

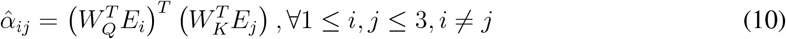

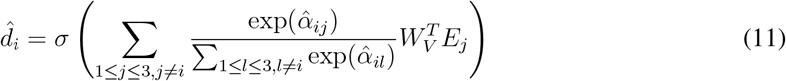

where 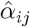 represent the attention coefficients before softmax normalization. 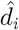 is calculated based on these coefficients as the weighted sum of linear transformed features with activation function.

We enhance the self-attention layer with the ReZero technique introduced in [52]. As noted in their study, ReZero accelerates model convergence and enables the use of additional stacked self-attention layers, improving the capacity and flexibility of the Hyper-SAGNN architecture. ReZero modifies the dynamic embedding calculation as: 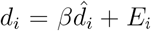, where *β* is a trainable parameter initialized as 0, as recommended by ReZero, and optimized during training.

Next, the Hadamard power (element-wise power) of the difference between each static *s*_*i*_ and dynamic (*d*_*i*_) embedding pair is calculated. This result is passed through a one-layer neural network with a linear activation function to produce a scalar value 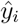. Finally, all 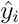 are averaged to reach the final predicted trigenic interaction score 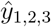:

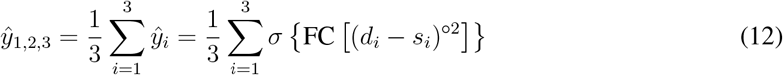

The hypergraph representation learning module is trained in an end-to-end manner with the log-cosh loss function:

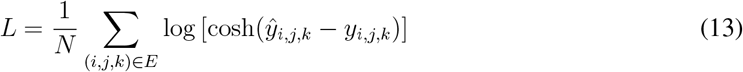

where *y*_*i*,*j*,*k*_ represents the true trigenic interaction score, and 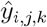 is the predicted score.

The log-cosh regression loss behaves like the MSE around zero and transitions to the mean absolute error (MAE) for larger target values, making it robust to outliers while maintaining stable gradients near zero [53].

### Uncertainty estimate using Gaussian process regression

To enhance explainability, we adopt a Gaussian process (GP) regression framework for uncertainty scoring [33, 34]. After training the Hyper-SAGNN model, we fit a GP model on the residuals between the predicted trigenic interaction score from Dango and the true score (*y*_*i*_) for each input trigenic interaction (*x*_*i*_).

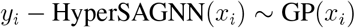

As demonstrated by [33], the trained GP model, fit on the residuals, can refine the original Dango predictions. By adding the mean of the GP to the original Dango score, we obtain an improved overall prediction 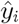 for the trigenic interaction:

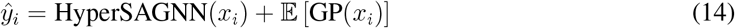

To quantify the model’s the uncertainty in predicting the genetic interaction *x*_*i*_, we calculated the standard deviation of the GP:

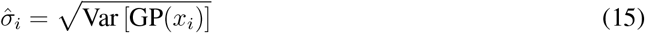

This predicted uncertainty score 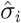 provides an interpretable measure of confidence for each predicted trigenic interaction, complementing the accuracy of the model’s predictions.

## Code Availability

Source code of Dango can be accessed at: https://github.com/ma-compbio/DANGO.

## Author Contributions

Conceptualization, R.Z., J-Z.M., and J.M.; Methodology, R.Z., M.B., J-Z.M., and J.M.; Software, R.Z., M.B.; Investigation, R.Z., M.B., J-Z.M., and J.M.; Writing – Original Draft, R.Z., J-Z.M., and J.M.; Writing – Review & Editing, R.Z., M.B., J-Z.M., and J.M.

## References

[1] Costanzo, M. et al. Global genetic networks and the genotype-to-phenotype relationship. Cell 177, 85–100 (2019).

[2] Mackay, T. F. & Anholt, R. R. Pleiotropy, epistasis and the genetic architecture of quantitative traits. Nature Reviews Genetics 1–19 (2024).

[3] Boone, C., Bussey, H. & Andrews, B. J. Exploring genetic interactions and networks with yeast. Nature Reviews Genetics 8, 437–449 (2007).

[4] Cowen, L., Ideker, T., Raphael, B. J. & Sharan, R. Network propagation: a universal amplifier of genetic associations. Nature Reviews Genetics 18, 551 (2017).

[5] Domingo, J., Baeza-Centurion, P. & Lehner, B. The causes and consequences of genetic interactions (epistasis). Annual Review of Genomics and Human Genetics 20, 433–460 (2019).

[6] Przybyla, L. & Gilbert, L. A. A new era in functional genomics screens. Nature Reviews Genetics 23, 89–103 (2022).

[7] Zitnik, M. et al. Current and future directions in network biology. Bioinformatics Advances 4, vbae099 (2024).

[8] Costanzo, M. et al. A global genetic interaction network maps a wiring diagram of cellular function. Science 353 (2016).

[9] Kuzmin, E. et al. Systematic analysis of complex genetic interactions. Science 360 (2018).

[10] Costanzo, M. et al. Environmental robustness of the global yeast genetic interaction network. Science 372, eabf8424 (2021).

[11] Ryan, C. J., Devakumar, L. P. S., Pettitt, S. J. & Lord, C. J. Complex synthetic lethality in cancer. Nature Genetics 55, 2039–2048 (2023).

[12] Beh, C. T., Cool, L., Phillips, J. & Rine, J. Overlapping functions of the yeast oxysterol-binding protein homologues. Genetics 157, 1117–1140 (2001).

[13] Suzuki, Y. et al. Knocking out multigene redundancies via cycles of sexual assortment and fluorescence selection. Nature Methods 8, 159–164 (2011).

[14] Bao, Z. et al. Homology-integrated CRISPR–Cas (HI-CRISPR) system for one-step multigene disruption in Saccharomyces cerevisiae. ACS Synthetic Biology 4, 585–594 (2015).

[15] Garst, A. D. et al. Genome-wide mapping of mutations at single-nucleotide resolution for protein, metabolic and genome engineering. Nature Biotechnology 35, 48–55 (2017).

[16] Lian, J., HamediRad, M., Hu, S. & Zhao, H. Combinatorial metabolic engineering using an orthogonal tri-functional CRISPR system. Nature Communications 8, 1–9 (2017).

[17] Zhang, H. et al. Simultaneous zygotic inactivation of multiple genes in mouse through CRISPR/Cas9-mediated base editing. Development 145 (2018).

[18] Zhang, Y. et al. A gRNA-tRNA array for CRISPR-Cas9 based rapid multiplexed genome editing in Saccharomyces cerevisiae. Nature Communications 10, 1–10 (2019).

[19] Celaj, A. et al. Highly combinatorial genetic interaction analysis reveals a multi-drug transporter influence network. Cell Systems 10, 25–38 (2020).

[20] Franceschini, A. et al. STRING v9.1: protein-protein interaction networks, with increased coverage and integration. Nucleic Acids Research 41, D808–D815 (2012).

[21] Zhang, R., Zou, Y. & Ma, J. Hyper-SAGNN: a self-attention based graph neural network for hypergraphs. In International Conference on Learning Representations (ICLR) (2020).

[22] Cho, H., Berger, B. & Peng, J. Compact integration of multi-network topology for functional analysis of genes. Cell Systems 3, 540–548 (2016).

[23] Eid, F.-E. et al. Systematic auditing is essential to debiasing machine learning in biology. Communications Biology 4, 183 (2021).

[24] Parenteau, J. et al. Deletion of many yeast introns reveals a minority of genes that require splicing for function. Molecular Biology of the Cell 19, 1932–1941 (2008).

[25] Feyder, S., De Craene, J.-O., Bär, S., Bertazzi, D. L. & Friant, S. Membrane trafficking in the yeast Saccharomyces cerevisiae model. International Journal of Molecular Sciences 16, 1509–1525 (2015).

[26] Sawin, K. E. & Tran, P. Cytoplasmic microtubule organization in fission yeast. Yeast 23, 1001–1014 (2006).

[27] Kanehisa, M. & Goto, S. KEGG: kyoto encyclopedia of genes and genomes. Nucleic Acids Research 28, 27–30 (2000).

[28] Stark, C. et al. BioGRID: a general repository for interaction datasets. Nucleic Acids Research 34, D535–D539 (2006).

[29] Pu, S., Wong, J., Turner, B., Cho, E. & Wodak, S. J. Up-to-date catalogues of yeast protein complexes. Nucleic Acids Research 37, 825–831 (2009).

[30] Peter, J. et al. Genome evolution across 1,011 Saccharomyces cerevisiae isolates. Nature 556, 339–344 (2018).

[31] Bepler, T. & Berger, B. Learning the protein language: Evolution, structure, and function. Cell Systems 12, 654–669 (2021).

[32] Lin, Z. et al. Evolutionary-scale prediction of atomic-level protein structure with a language model. Science 379, 1123–1130 (2023).

[33] Qiu, X., Meyerson, E. & Miikkulainen, R. Quantifying point-prediction uncertainty in neural networks via residual estimation with an i/o kernel. arXiv preprint 1906.00588 (2019).

[34] Hie, B., Bryson, B. D. & Berger, B. Leveraging uncertainty in machine learning accelerates biological discovery and design. Cell Systems 11, 461–477 (2020).

[35] Sterner, D. E. et al. Functional organization of the yeast SAGA complex: distinct components involved in structural integrity, nucleosome acetylation, and TATA-binding protein interaction. Molecular and Cellular Biology 19, 86–98 (1999).

[36] Liu, G. et al. Architecture of Saccharomyces cerevisiae SAGA complex. Cell Discovery 5, 25 (2019).

[37] Pray-Grant, M. G. et al. The novel SLIK histone acetyltransferase complex functions in the yeast retrograde response pathway. Molecular and Cellular Biology 22, 8774–8786 (2002).

[38] Rood, J. E., Hupalowska, A. & Regev, A. Toward a foundation model of causal cell and tissue biology with a perturbation cell and tissue atlas. Cell 187, 4520–4545 (2024).

[39] Roohani, Y., Huang, K. & Leskovec, J. Predicting transcriptional outcomes of novel multigene perturbations with GEARS. Nature Biotechnology 42, 927–935 (2024).

[40] Han, K. et al. Synergistic drug combinations for cancer identified in a CRISPR screen for pairwise genetic interactions. Nature Biotechnology 35, 463–474 (2017).

[41] Berg, J. & Lässig, M. Cross-species analysis of biological networks by bayesian alignment. Proceedings of the National Academy of Sciences 103, 10967–10972 (2006).

[42] Hirsh, E. & Sharan, R. Identification of conserved protein complexes based on a model of protein network evolution. Bioinformatics 23, e170–e176 (2007).

[43] Zhou, D., Huang, J. & Schölkopf, B. Learning with hypergraphs: Clustering, classification, and embedding. In Advances in Neural Information Processing Systems, 1601–1608 (2007).

[44] Tu, K., Cui, P., Wang, X., Wang, F. & Zhu, W. Structural deep embedding for hyper-networks. In Thirty-Second AAAI Conference on Artificial Intelligence (2018).

[45] Gao, Y., Feng, Y., Ji, S. & Ji, R. HGNN+: General hypergraph neural networks. IEEE Transactions on Pattern Analysis and Machine Intelligence 45, 3181–3199 (2022).

[46] Li, W. et al. scMHNN: a novel hypergraph neural network for integrative analysis of single-cell epigenomic, transcriptomic and proteomic data. Briefings in Bioinformatics 24, bbad391 (2023).

[47] Hamilton, W., Ying, Z. & Leskovec, J. Inductive representation learning on large graphs. In Advances in Neural Information Processing Systems, 1024–1034 (2017).

[48] Kiela, D., Wang, C. & Cho, K. Dynamic meta-embeddings for improved sentence representations. In Proceedings of the 2018 Conference on Empirical Methods in Natural Language Processing (EMNLP) (Brussels, Belgium, 2018).

[49] Xie, Y., Hu, Y., Xing, L. & Wei, X. Dynamic task-specific factors for meta-embedding. In International Conference on Knowledge Science, Engineering and Management, 63–74 (Springer, 2019).

[50] Liu, Q. et al. Domain-specific meta-embedding with latent semantic structures. Information Sciences (2020).

[51] Vaswani, A. et al. Attention is all you need. In Advances in Neural Information Processing Systems, 5998–6008 (2017).

[52] Bachlechner, T., Majumder, B. P., Mao, H. H., Cottrell, G. W. & McAuley, J. Rezero is all you need: Fast convergence at large depth. arXiv preprint 2003.04887 (2020).

[53] Neuneier, R. & Zimmermann, H. G. How to train neural networks. In Neural networks: tricks of the trade, 373–423 (Springer, 1998).

