## Supplemental Information for "DANGO: Predicting higher-order genetic interactions"

#### A Supplementary Methods

##### A.1 Choice of hypergraph representation learning framework for DANGO

We use Hyper-SAGNN in DANGO over other graph representation learning methods for several important reasons. First, the hypergraph in this work is a homogeneous hypergraph (i.e., no intrinsic order in each hyperedge). In other words, to make predictions on the hyperedges, the model must be permutation invariant to the order of input. Second, the model needs to handle non-fixed-size input to predict higher-order interactions beyond trigenic interactions. Most importantly, as defined in the main text, trigenic interactions are characterized as the additional synergy beyond the expected phenotypes from lower-order interactions. This requires that hyperedges corresponding to trigenic interactions be indecomposable, meaning decomposition of the hyperedge would result in the loss of higher-order information.

Considering these requirements, most existing graph representation learning methods for pairwise interactions [1, 2] and hyperedge prediction methods [3, 4] are unsuitable because they cannot capture the higher-order information within indecomposable hyperedges or require fixed-sized and ordered input. For each gene $A$ in gene set $S$, its node representation depends on the difference between two types of embeddings: one based on its features and the other one determined by the features of the remaining genes $S \setminus A$. As a result, there are no features solely dependent on one node, and the final prediction is also indecomposable. Our Hyper-SAGNN [5] is applicable to both homogeneous and heterogeneous hypergraphs with variable hyperedge sizes and achieves state-of-the-art performances in multiple applications [5, 6]. In this work, we further enhance Hyper-SAGNN with pre-trained GNNs and a meta embedding learning module, enabling it to incorporate interaction information from various sources.

##### A.2 Training procedure of DANGO

During training, all components of DANGO, including the six pretrained GNNs, meta embedding learning module, and the modified Hyper-SAGNN, are jointly optimized in an end-to-end manner using the log-cosh regression loss described in Eqn. 13 in the main text. Neural networks are trained using the Adam optimizer [7] with a learning rate of 1e-3. Training terminates when performance on a held-out validation set does not improve for 20 consecutive epochs. The best model parameters, determined based on the validation set, are used for test set evaluation or *de novo* predictions.

To improve prediction robustness, we employ an ensemble technique by training multiple models with identical structures but different random initialization. Final predictions are averaged over these models. Experiments show that increasing the number of models enhance performance stability (see **Fig.** S1). For efficiency, the ensemble size is set to 10 to balance performance and training time. For 5-fold cross-validation, 20% of the data is assigned to the test set in each fold, while the remaining 80% is randomly split into training and validation sets in a 4:1 ratio. Models are trained ten times within each fold, ensuring robust evaluation.

***Computational complexity***: All evaluations were performed on a 32-core machine with an NVIDIA GeForce 2080Ti GPU. The 5-fold cross-validation process, involving training 10 DANGO models per fold, took approximately 12 hours. Predictions for set #2, containing around 451 million triplets, took 8 hours.

##### A.3 Design of the baseline models

We adapted existing algorithms for pairwise interaction prediction to trigenic interaction prediction to serve as baseline methods. Three baseline methods were designed for comparison. The first two models use learned gene embedding from the six PPI networks in the STRING database (v9.1) generated using the Mashup method [8]. These embeddings are averaged for each triplet and input into decision tree ensemble methods: random forest and gradient boosting decision tree. These methods are referred to as Mashup-RF and Mashup-GTB, respectively. To ensure fair comparisons, we also included a baseline trained in an end-to-end manner similar to DANGO, referred to as GCN-Avg-DNN. In this model, gene embeddings are generated through six pre-trained GNNS as in DANGO. However, instead of the meta embedding learn module, the six embedding vectors for each node are averaged to create the final node embedding. These averaged embeddings are further averaged for each tripet and input into a deep neural network regression model.

##### A.4 Enforcing trigenic interactions in the neural network structure

We implemented a two-layered DNN for predicting yeast growth, with input size equal to the number of considered SNPs $S$ and output size equal to the number of conditions (see **Fig.** 5a. The hidden layer size ($E$) matches the number of selected trigenic interactions, which was set to 1,395. To cover a broad range of genes, we selected trigenic interactions such that each gene appears in at least one interaction, prioritizing those with the highest predicted scores.

The first layer of the DNN is sparsely connected, with a weight matrix $W$ of size $E \times S$. The structure of the trigenic interactions is enforced by setting only relevant entries in $W$ to non-zero values. For example, if a trigenic interaction $E_i$ involves genes $(\text{gene}_1, \text{gene}_2, \text{gene}_3)$, and $\text{SNP}_{1-2}, \text{SNP}_{3-4}, \text{SNP}_5$ reside in these genes, respectively, only the corresponding columns (1st to 5th columns) in the $i$-th row of $W$ are set to non-zero. Alternatively, SNPs can be grouped by genes before being used as features, reducing $W$ to size $E \times G$, where $G$ is the number of genes.

#### References


[1] Perozzi, B., Al-Rfou, R. & Skiena, S. Deepwalk: Online learning of social representations. In *Proceedings of the 20th ACM SIGKDD international conference on Knowledge Discovery and Data Mining*, 701–710 (ACM, 2014).

[2] Grover, A. & Leskovec, J. node2vec: Scalable feature learning for networks. In *Proceedings of the 22nd ACM SIGKDD International Conference on Knowledge Discovery and Data Mining*, 855–864 (ACM, 2016).

[3] Tu, K., Cui, P., Wang, X., Wang, F. & Zhu, W. Structural deep embedding for hyper-networks. In *Thirty-Second AAAI Conference on Artificial Intelligence* (2018).

[4] Gui, H. *et al.* Large-scale embedding learning in heterogeneous event data. In *2016 IEEE 16th International Conference on Data Mining (ICDM)*, 907–912 (IEEE, 2016).

[5] Zhang, R., Zou, Y. & Ma, J. Hyper-SAGNN: a self-attention based graph neural network for hypergraphs. In *International Conference on Learning Representations (ICLR)* (2020).

[6] Zhang, R. & Ma, J. MATCHA: Probing multi-way chromatin interaction with hypergraph representation learning. *Cell Systems* **10**, 397–407 (2020).

[7] Kingma, D. P. & Ba, J. Adam: A method for stochastic optimization. *arXiv preprint arXiv:1412.6980* (2014).

[8] Cho, H., Berger, B. & Peng, J. Compact integration of multi-network topology for functional analysis of genes. *Cell Systems* **3**, 540–548 (2016).

### B Supplementary Figures

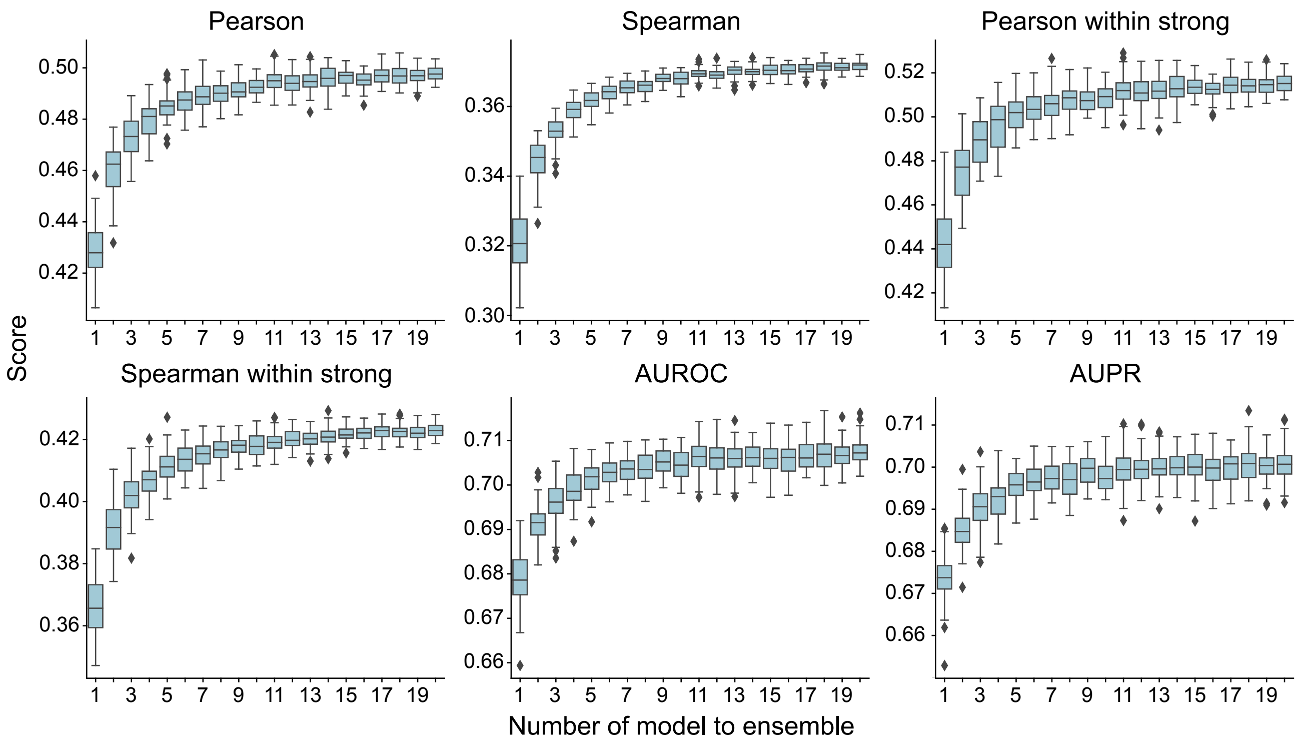


**Figure S1:** Performance evaluation for DANGO with different numbers of ensembled models.

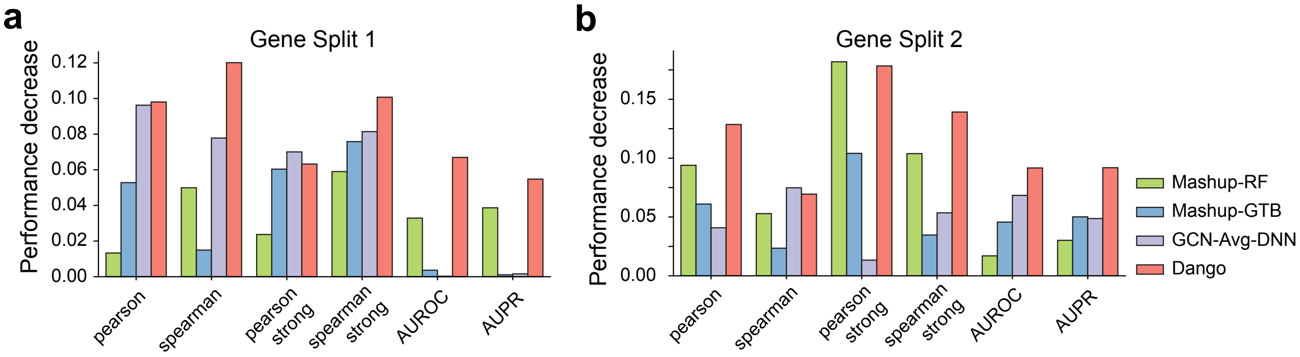


**Figure S2:** Performance decreases when using randomly shuffled node features. **a**. Results under gene split 1, where a fraction of genes in the test set are unobserved in the training dataset. **b**. Results under gene split 2, where all genes in the test set are unobserved in the training dataset.

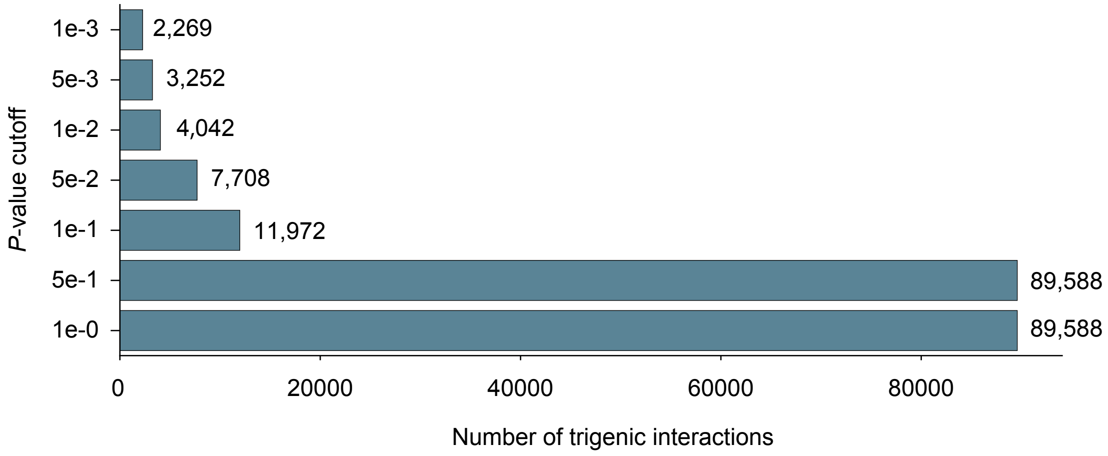


**Figure S3:** Number of measured trigenic interactions under different *P*-value cut-offs.

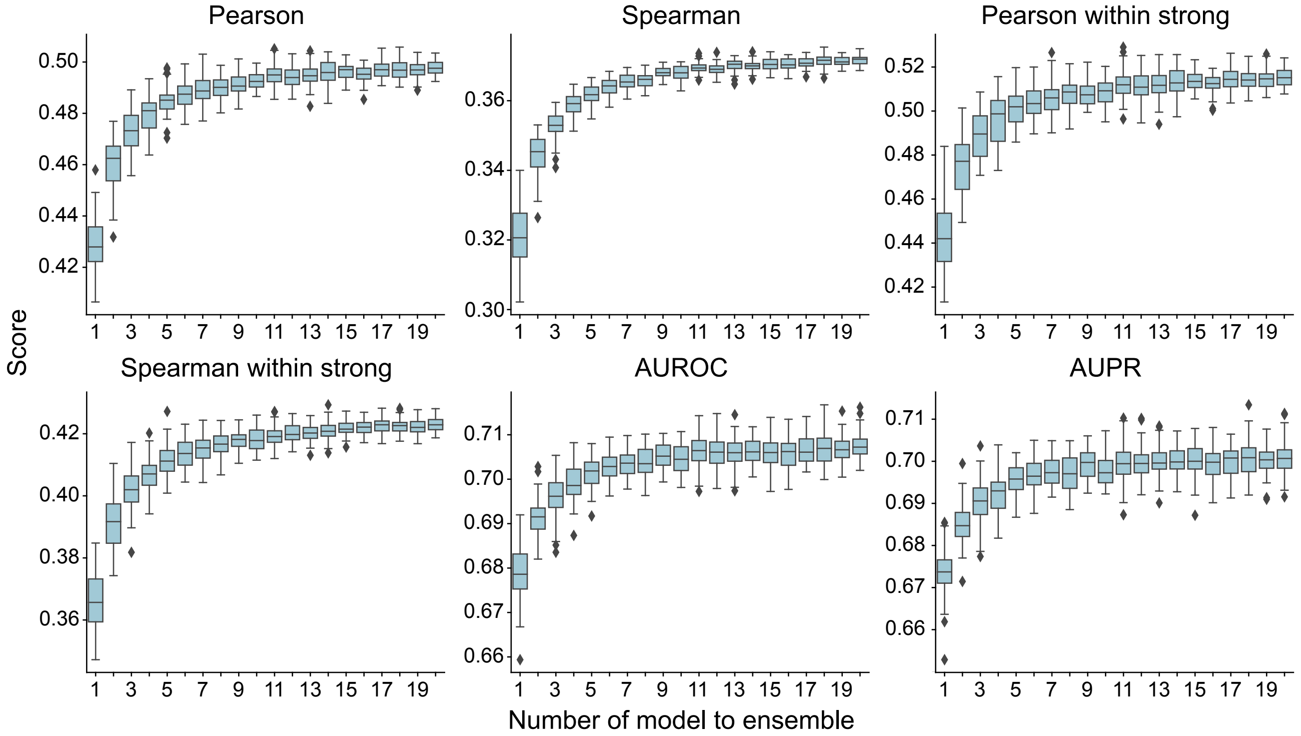


**Figure S4:** Performance evaluation for DANGO with different numbers of ensemble models.

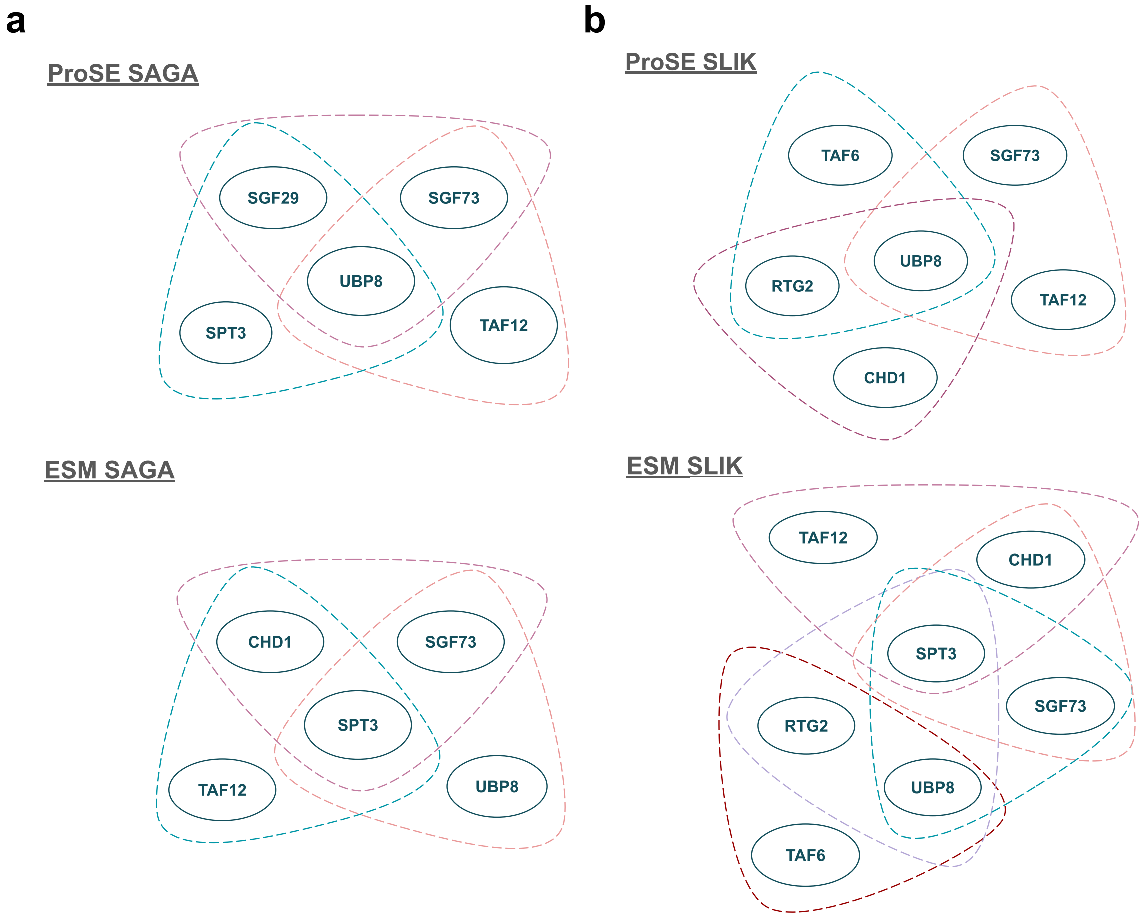


**Figure S5:** Visualization of predicted ProSE/ESM2-informed trigenic interactions. **a**. Predicted trigenic interactions associated with the SAGA yeast complex, using ProSE and ESM2 embeddings. **b**. Predicted trigenic interactions associated with the SAGA-like SLIK yeast complex, using ProSE and ESM2 embeddings)
